# NGFR/Ngfr-marked basal duct progenitors drive ductal–acinar regeneration in injured salivary glands

**DOI:** 10.64898/2026.05.13.724951

**Authors:** Seong Gyeong Jeon, Dong Hyuck Bae, Jun-Yeol Park, Manjung Yong, Thi Viet Trinh Nguyen, Kyung Jin Lee, Sang-Hyuk Lee, Young Chang Lim, Eun Ju Bae, Mi-Young Son, Jongman Yoo

## Abstract

Severe salivary gland injury can cause chronic xerostomia and persistent secretory dysfunction, yet the epithelial populations that support repair remain poorly defined. Here, we identify NGFR/Ngfr as a conserved surface marker for isolating organoid-forming epithelial stem/progenitor cells from human and mouse salivary glands and show that mouse Ngfr-lineage cells contribute to ductal–acinar regeneration after injury. Single-cell transcriptomic analysis of human salivary gland tissue identified a restricted NGFR-expressing basal duct epithelial subpopulation with progenitor-like features and early positions along inferred epithelial differentiation trajectories. Functionally, NGFR-expressing cells showed enhanced primary and secondary organoid-forming capacity, and NGFR-enriched human organoids engrafted after transplantation into injured salivary glands of immunodeficient mice. In mouse salivary glands, isolated Ngfr-expressing cells showed enriched organoid-forming activity, and Ngfr expression localized to injury-associated ductal regions after duct ligation and local inflammatory injury. Ngfr-CreERT2 lineage tracing further showed that Ngfr-lineage cells contribute to ductal and acinar compartments during post-injury regeneration. Together, these findings establish NGFR/Ngfr as a conserved surface marker for prospectively isolating basal duct epithelial stem/progenitor populations with organoid-forming activity and injury-responsive ductal–acinar regenerative potential.

## Introduction

The salivary gland epithelium is composed of acinar, ductal, and myoepithelial compartments that coordinate saliva production, modification, and expulsion (Barrows et al., 2020; Chibly et al., 2022). Acinar cells produce and secrete primary saliva, whereas ductal cells transport this fluid through the ductal network and modify its ionic composition. Myoepithelial cells, which surround acini and portions of the intercalated ducts, contract in response to autonomic stimulation to facilitate saliva expulsion and ductal flow (Barrows et al., 2020; Amano et al., 2025). The coordinated function of these epithelial compartments is essential for maintaining salivary gland physiology.

Radiotherapy for head and neck cancer frequently damages normal salivary gland tissue adjacent to the tumor, resulting in long-term salivary gland dysfunction. Functional impairment and loss of acinar cells are major contributors to hyposalivation, an objective reduction in salivary output, and xerostomia, the subjective sensation of oral dryness (May et al., 2018; Li et al., 2022). However, radiation-induced salivary gland dysfunction is not driven solely by acinar cell loss. Rather, it reflects a broader regenerative failure involving injury to stem/progenitor cell pools, ductal remodeling, disrupted neural and vascular support, stromal niche alterations, and fibrosis (May et al., 2018; Li et al., 2022; Grundmann et al., 2009). Current clinical treatments can transiently support residual gland function or relieve symptoms, but they do not rebuild the damaged salivary epithelium or restore long-term secretory capacity (Dirix et al., 2006; Chambers et al., 2007). Experimental regenerative approaches, including stem/progenitor cell and organoid-based strategies, have highlighted the therapeutic potential of epithelial replacement but also underscore the need to define repair-competent epithelial populations more precisely (Nanduri et al., 2011; Pringle et al., 2016; Rocchi et al., 2021; Jeon et al., 2024).

Unlike tissues maintained by a well-defined stem cell hierarchy, such as the intestinal epithelium or the hematopoietic system, the adult salivary gland epithelium is thought to be maintained largely through lineage-restricted mechanisms within individual epithelial compartments. Lineage tracing studies have shown that acinar cells are maintained primarily by self-duplication of pre-existing acinar cells rather than by continuous replenishment from a broadly multipotent stem cell pool (Aure et al., 2015). In the postnatal salivary gland, acinar, ductal, and myoepithelial lineages also exhibit relatively separated lineage relationships compared with earlier developmental stages (Aure et al., 2019). These findings support a model in which adult salivary gland homeostasis is governed not by a single dominant multipotent stem cell population, but by compartment-specific progenitors and self-renewal of differentiated epithelial cells. However, it remains unclear which epithelial cell population acquires injury-responsive properties and contributes to post-injury repair.

Translating injury-responsive epithelial populations into regenerative strategies requires their precise definition, prospective isolation, and reliable in vitro expansion (Rocchi et al., 2021; Maimets et al., 2016). Previous studies have proposed ductal progenitor-associated markers, including KIT and KRT5/KRT14 (May et al., 2018; Kwak et al., 2018), acinar or secretory progenitor-associated markers, including SOX2 and SOX9 (Emmerson et al., 2018; Xu et al., 2022), and surface markers used to enrich salivary epithelial or organoid-forming fractions, such as EpCAM, CD24, CD49f/ITGA6, and CD26/DPP4 (Maimets et al., 2016; Yoon et al., 2022; Aalam et al., 2025). Together, these studies have provided important tools for investigating salivary gland epithelial heterogeneity and regenerative potential. However, a human tissue-validated surface marker that is compatible with live-cell isolation and sufficiently specific to distinguish a defined repair-associated epithelial population from neighboring epithelial compartments remains needed.

NGFR, also known as p75NTR or CD271, was originally characterized as a neurotrophin receptor but has also been implicated as a surface marker associated with stem/progenitor-enriched basal epithelial states in several tissues, including the esophageal epithelium, oral mucosa, corneal limbus, and airway epithelium (Okumura et al., 2003; Nakamura et al., 2007; Di Girolamo et al., 2008; Rock et al., 2009). However, whether NGFR marks a defined basal duct epithelial population with repair-associated properties in the salivary gland remains unknown. In this study, we combine human salivary gland single-cell transcriptomics, prospective cell isolation, organoid-based functional assays, transplantation, and mouse injury lineage tracing to identify NGFR/Ngfr as a conserved marker of an injury-responsive epithelial population involved in salivary gland regeneration.

## Results

### NGFR marks a restricted basal duct epithelial subpopulation distinct from the myoepithelial compartment in the human salivary gland

In a previous study, we established a human salivary gland organoid culture system and demonstrated the regenerative potential of these organoids (Jeon et al., 2024). However, the endogenous epithelial population enriched for organoid formation remained undefined. To identify candidate surface markers for the isolation of organoid-forming cells, we performed single-cell RNA sequencing on two normal human submandibular gland tissue samples (Sample 1 and Sample 2), yielding 11,161 high-quality cells after quality control filtering (Supplementary Fig. 1a, b). Unsupervised clustering and UMAP visualization identified 17 cell populations, which were annotated based on canonical marker gene expression (Fig. 1a). These included epithelial populations, such as acinar, mucous acinar, ductal, basal, and myoepithelial cells, as well as immune, endothelial, and stromal cell populations. The distribution of annotated cell types in each sample is shown in Supplementary Fig. 1c. Cell identities were supported by expression of established markers, including KRT5 and KRT14 for basal epithelial cells, AQP5 and PRB3 for acinar cells, KRT19 and KRT7 for ductal cells, ACTA2 and MYLK for myoepithelial cells, CD3E for T cells, IGHM for B cells, and PECAM1 and VWF for endothelial cells (Fig. 1b). Representative marker gene expression supporting the annotation of all 17 clusters is shown in a heatmap (Supplementary Fig. 1e). To characterize epithelial heterogeneity in greater detail, epithelial cells were extracted and re-clustered, revealing five major epithelial subpopulations: acinar, ductal, basal, mucous acinar, and myoepithelial cells (Fig. 1c). The basal epithelial cluster was transcriptionally distinct from differentiated acinar and ductal populations and was enriched for basal epithelial markers, including KRT5 and TP63 (Fig. 1d). Because our previous work suggested that basal duct-associated cells may contain organoid-forming potential, we focused on identifying surface markers enriched within this compartment.

**Figure 1.**
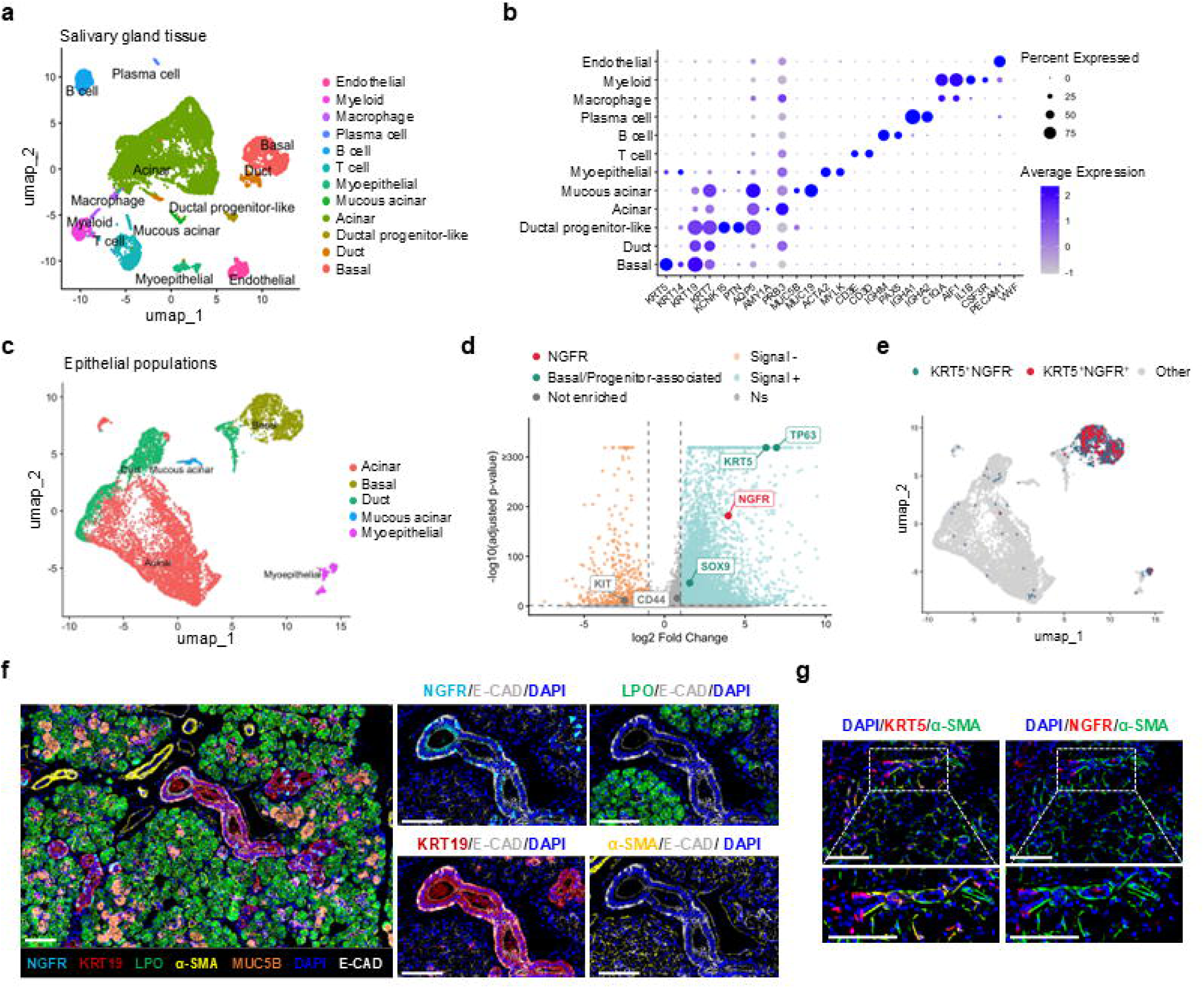
NGFR marks a restricted basal duct-associated epithelial subpopulation distinct from the myoepithelial compartment in the human salivary gland. **a** UMAP visualization of 11,161 cells from two normal submandibular gland tissue samples, colored by annotated cell type. **b** Dot plot showing expression of canonical marker genes across cell types. Dot size indicates the percentage of expressing cells; color intensity indicates average expression level. **c** UMAP of epithelial subset after re-clustering, revealing five distinct epithelial subpopulations: Acinar, Duct, Basal, Mucous acinar, and Myoepithelial. **d** Volcano plot of differentially expressed genes in the basal cluster versus other epithelial cells. Stem/progenitor-associated genes KRT5, TP63, and SOX9 are highlighted in green, whereas KIT and CD44, which showed low enrichment in the salivary gland basal cluster, are shown in gray. NGFR is highlighted in red (log_2_FC = 3.95, adjusted p-value < 2.4×10^-182^). e. UMAP of epithelial cells colored by KRT5/NGFR co-expression status. KRT5^+^NGFR^+^ (Red) and KRT5^+^NGFR^-^ (Blue) cells are shown. KRT5 is expressed in 77.9% of basal cells, whereas NGFR expression is restricted to 16.9%, with KRT5^+^NGFR^+^ cells comprising 15.7% of the basal population. f. Multiplex immunofluorescence staining of human salivary gland tissue for NGFR (cyan), KRT19 (red), LPO (green), α-SMA (yellow), MUC5B (orange), and E-CAD (white). Scale bar, 100 μm. g. Spatial distribution of KRT5, NGFR, and α-SMA with minimal overlap between NGFR and α-SMA. Scale bar, 100 μm.

Differential expression analysis of the basal cluster identified NGFR as one of the most strongly enriched cell-surface receptor genes (log2FC = 3.95; adjusted p-value < 2.4 × 10^−182^). Given its strong enrichment and compatibility with live-cell isolation, we selected NGFR as a candidate surface marker for further validation (Fig. 1d). We next assessed the relationship between NGFR and the broader basal marker KRT5. KRT5 was expressed in 77.9% of basal cells, whereas NGFR was restricted to 16.9% of basal cells. KRT5^+^NGFR^+^ cells accounted for 15.7% of the basal population, and 93% of NGFR^+^ cells overlapped with KRT5^+^ cells. NGFR^+^ cells were detected in both tissue samples (Sample 1: 0.79%; Sample 2: 4.15% of total cells; Supplementary Fig. 1d). Thus, although most NGFR^+^ cells were contained within the KRT5^+^ basal compartment, NGFR marked a substantially narrower basal epithelial subpopulation than KRT5 (Fig. 1e).

We next performed multiplex immunostaining to validate the spatial localization of NGFR^+^ cells in human salivary gland tissue. Consistent with the scRNA-seq analysis, NGFR was restricted to a subset of basal epithelial cells associated with ductal structures and showed minimal overlap with acinar or myoepithelial cells, localizing primarily within the ductal compartment (Fig. 1f). In contrast, KRT5 broadly labeled both basal ductal cells and myoepithelial cells, whereas NGFR more selectively marked a basal duct-associated epithelial subpopulation distinct from the myoepithelial compartment (Fig. 1g). Together, these findings identify NGFR as a candidate surface marker for a restricted basal duct-associated epithelial subpopulation distinct from the myoepithelial compartment in the human salivary gland.

### NGFR-expressing basal ductal cells exhibit a progenitor-like transcriptional state within the salivary gland epithelium

To quantitatively assess the relative differentiation state of salivary gland epithelial cells, we performed CytoTRACE2 analysis. Projection of CytoTRACE2 scores on the epithelial UMAP revealed that the Basal cluster exhibited markedly higher scores compared to differentiated epithelial populations including Acinar, Duct, and Mucous acinar cells (Fig. 2a). Comparison of scores across epithelial subtypes demonstrated that the Basal cluster had the highest mean score (0.251), followed by Duct (0.094), Mucous acinar (0.079), and Acinar (0.068), consistent with the expected differentiation hierarchy (Fig. 2b). Potency category analysis showed that while Acinar, Duct, and Mucous acinar clusters were predominantly classified as Differentiated, the Basal cluster contained a significantly higher proportion of Oligopotent and Unipotent cells, suggesting that basal epithelial cells retain a less differentiated transcriptional state compared with other epithelial populations (Fig. 2c, left).

**Figure 2.**
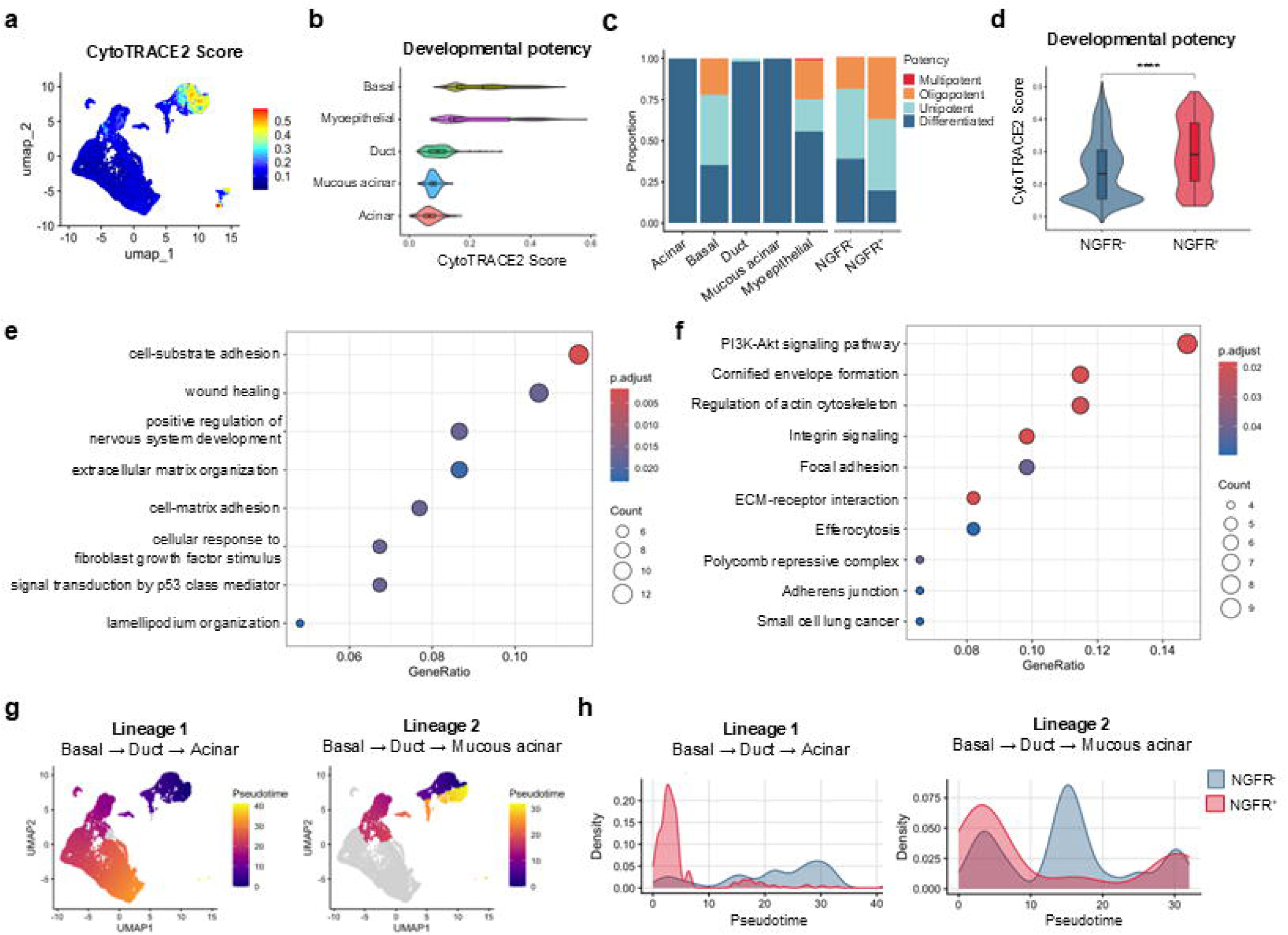
NGFR-positive basal cells occupy an early progenitor-like transcriptional state in the salivary gland epithelium. a. UMAP visualization of CytoTRACE2 developmental potency scores across epithelial cell populations. Higher scores indicate less differentiated states. b. Violin plots comparing CytoTRACE2 scores across epithelial subtypes. The basal cluster exhibits the highest CytoTRACE2 scores. c. Stacked bar plots showing the distribution of CytoTRACE2 potency categories (Multipotent, Oligopotent, Unipotent, and Differentiated) across epithelial subtypes (Left) and between NGFR^+^ and NGFR^-^ cells within the basal cluster (Right). NGFR^+^ cells are enriched for the Oligopotent category relative to NGFR^−^ cells. d. Violin plots comparing CytoTRACE2 scores between NGFR^+^ and NGFR^-^ cells within the Basal cluster. Statistical significance was determined by Wilcoxon rank-sum test (****p < 0.0001). e. Dot plot of Gene Ontology Biological Process enrichment analysis of genes significantly upregulated in NGFR^+^ cells. Dot size indicates gene count; color indicates adjusted p value. f. Dot plot of KEGG pathway enrichment analysis of genes significantly upregulated in NGFR^+^ cells. Enriched pathways include PI3K-Akt signaling, integrin signaling, focal adhesion, and ECM-receptor interaction. Dot size indicates gene count, and color indicates adjusted p value. g. UMAP visualization of pseudotime values along two differentiation lineages inferred by Slingshot trajectory analysis. Lineage 1: Basal → Duct → Acinar; Lineage 2: Basal → Duct → Mucous acinar. h. Density plots showing the pseudotime distribution of NGFR^+^ and NGFR^-^ cells along Lineage 1 (Left) and Lineage 2 (Right). NGFR^+^ cells are concentrated at early pseudotime values in both lineages.

Comparison of CytoTRACE2-inferred potency between NGFR^+^ and NGFR^-^ cells within the Basal cluster revealed that NGFR^+^ cells exhibited significantly higher CytoTRACE2 scores than NGFR- cells (NGFR^+^: 0.294 vs NGFR^-^: 0.243, p < 0.0001, Fig. 2d). Consistently, potency category analysis demonstrated that the proportion of Oligopotent cells was approximately two-fold higher in NGFR^+^ cells compared to NGFR^-^ cells (35.2% vs 19.1%), indicating that NGFR^+^ cells are enriched for a progenitor-like transcriptional state within the Basal cluster (Fig. 2c, right). Gene ontology analysis revealed that genes upregulated in NGFR^+^ cells were significantly enriched for biological processes including cell-substrate adhesion (adjusted p value = 0.0017), wound healing, extracellular matrix organization, cell-matrix adhesion, and cellular response to fibroblast growth factor stimulus (Fig. 2e). Consistent with these findings, KEGG pathway analysis identified significant enrichment of PI3K-Akt signaling, integrin signaling, focal adhesion, and ECM-receptor interaction pathways, further supporting a progenitor-associated transcriptional program in NGFR^+^ cells (Fig. 2f).

Slingshot trajectory analysis was performed to infer potential epithelial differentiation trajectories in the salivary gland. The Basal cluster was identified as the common origin of two distinct differentiation lineages: Lineage 1 (Basal → Duct → Acinar) and Lineage 2 (Basal → Duct → Mucous acinar) (Fig. 2g). Pseudotime distribution analysis demonstrated that NGFR^+^ cells were concentrated at early pseudotime values in both lineages, whereas NGFR^-^ cells were broadly distributed across the entire pseudotime range (Fig. 2h). These findings suggest that NGFR^+^ basal cells occupy an early position within inferred ductal–acinar and ductal–mucous acinar epithelial trajectories, consistent with a progenitor-like transcriptional state rather than a fully differentiated epithelial identity.

### NGFR-based isolation prospectively enriches organoid-forming epithelial cells from human salivary gland tissue

To test whether NGFR enriches cells with organoid-forming activity, we isolated NGFR^+^ and NGFR^−^ cells from human salivary gland tissue and compared their organoid-forming efficiency (Fig. 3a). NGFR^+^ cells represented approximately 5% of the epithelial population before sorting (Fig. 3b). After 7 days of culture, the NGFR^-^ fraction exhibited an organoid-forming efficiency of approximately 0.2%, whereas the NGFR^+^ fraction showed an organoid-forming efficiency of approximately 6.5% (Fig. 3c). These results indicate that organoid-forming activity is markedly enriched within the NGFR^+^ fraction compared with the NGFR^-^ fraction.

**Figure 3.**
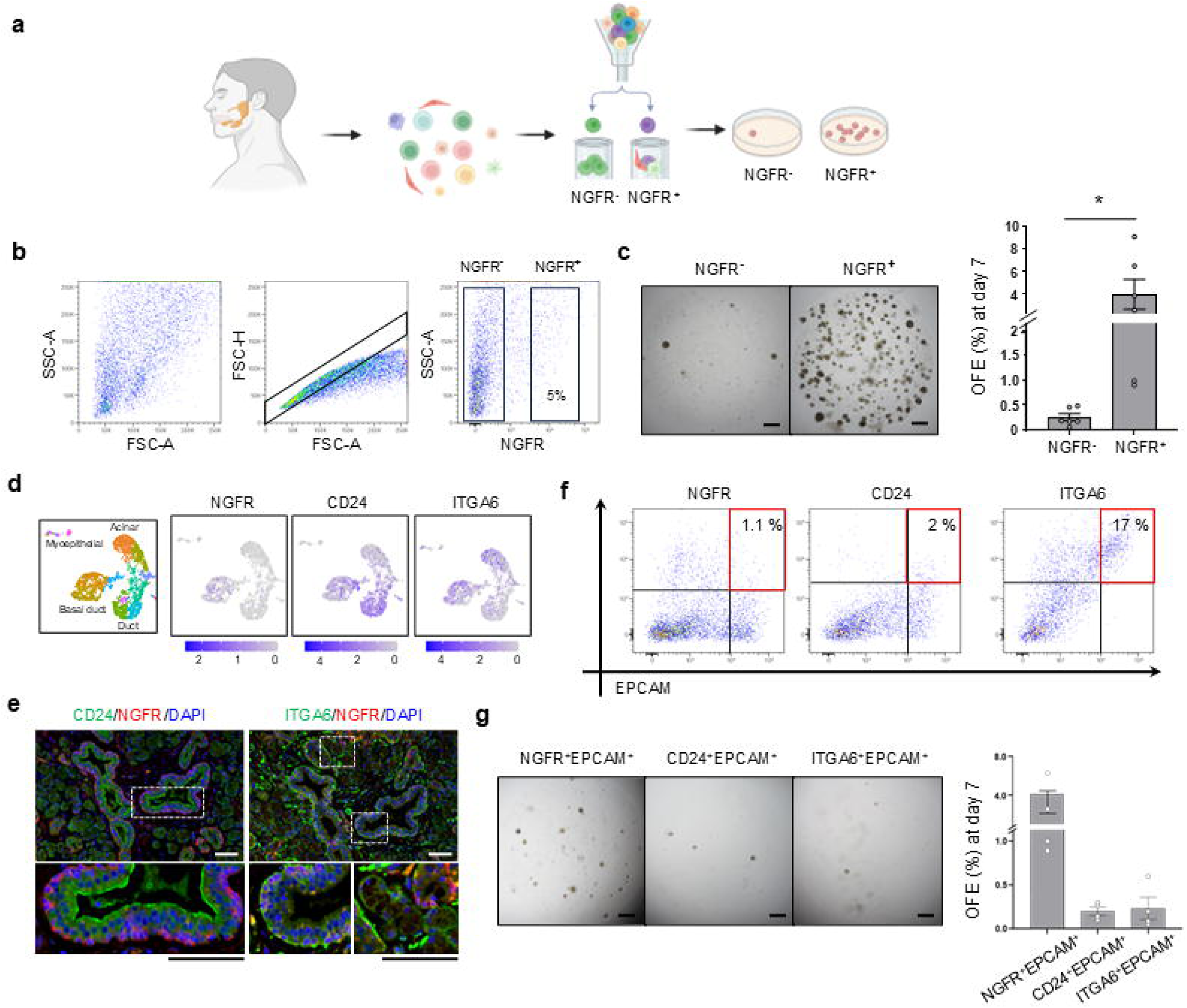
NGFR enriches for human salivary gland epithelial cells with high organoid-forming capacity. a. Schematic of NGFR-based prospective isolation of human salivary gland cells and subsequent organoid-forming assay. Dissociated human salivary gland cells were sorted into NGFR^+^ and NGFR^−^ fractions by FACS and cultured in parallel to compare organoid-forming capacity. b. Flow cytometric gating strategy for dissociated human salivary gland cells, including all events, singlet discrimination, and identification of NGFR^-^ and NGFR^+^ populations. NGFR⁺ cells accounted for approximately 5% of the dissociated cell population. c. Bright-field images and quantification of organoid-forming efficiency (OFE) of sorted NGFR^-^ and NGFR^+^ cells from human salivary gland tissue. *p < 0.05. Scale bar, 500 μm. d. Feature plots of the distribution of NGFR, CD24, and ITGA6 expression in single-cell RNA-sequencing data from human salivary gland tissue. e. Immunofluorescence staining of NGFR with CD24 or ITGA6 in human salivary gland. Higher-magnification views of the boxed regions are shown on the bottom. Scale bar, 50 μm. f. FACS plots showing the isolation of NGFR^+^EPCAM^+^, CD24^+^EPCAM^+^, and ITGA6^+^EPCAM^+^ populations from dissociated human salivary gland tissue. g. Bright-field images and quantification of organoid-forming efficiency of sorted NGFR^+^EPCAM^+^, CD24^+^EPCAM^+^, and ITGA6^+^EPCAM^+^ cells. Data are shown as mean ± SEM. Scale bar, 500 μm.

We next compared the functional utility of NGFR with previously proposed candidate markers, including CD24 and ITGA6/CD49f, which have been used to isolate salivary gland stem/progenitor-enriched epithelial fractions (Aalam et al., 2025; Yoon et al., 2022). Single-cell RNA sequencing analysis showed that NGFR, CD24, and ITGA6 were enriched in partially distinct epithelial compartments (Fig. 3d). Immunostaining confirmed that CD24 and ITGA6 marked epithelial compartments distinct from NGFR. CD24 was predominantly expressed along the luminal surface of ductal epithelium with minimal overlap with NGFR^+^ basal ductal cells, whereas ITGA6 was broadly distributed around basement membrane-associated epithelial regions, including basal ductal areas (Fig. 3e). These findings indicate that NGFR identifies a more discrete basal duct-associated epithelial subpopulation than CD24 or ITGA6. To directly compare their ability to enrich organoid-forming activity, NGFR^+^EPCAM^+^, CD24^+^EPCAM^+^, and ITGA6^+^EPCAM^+^ fractions were isolated and cultured under identical conditions. The NGFR^+^EPCAM^+^ fraction exhibited the highest organoid-forming efficiency, exceeding that of the CD24^+^EPCAM^+^ and ITGA6^+^EPCAM^+^ fractions (Fig. 3f, g). These findings suggest that NGFR more efficiently enriches organoid-forming activity than other candidate markers tested. Together, these results show that NGFR^+^ cells constitute a relatively rare population within human salivary gland tissue, yet the NGFR^+^ fraction is strongly enriched for organoid-forming capacity.

### NGFR expression defines a progenitor-enriched organoid state with enhanced secondary organoid-forming capacity

Given that tissue-derived NGFR^+^ cells showed high organoid-forming efficiency, we next asked whether NGFR could also distinguish functionally distinct cell states within the organoid culture system. Analysis of NGFR expression in this system showed that more than 95% of cells expressed NGFR at the early culture stage. As culture progressed and organoids became more structurally organized, NGFR expression gradually became restricted to the outer layer of the organoids (Fig. 4a). We next examined the relationship between NGFR expression and proliferative status by assessing co-expression with Ki67. Although the proportion of NGFR^+^Ki67^+^ cells among total cells decreased from day 5 to day 15 of culture, the proportion of Ki67^+^ proliferating cells that retained NGFR expression remained high throughout the culture period (Fig. 4a). In parallel, expression of acinar differentiation markers, including AQP5 and AMY1A, increased over time, whereas NGFR expression declined together with the basal progenitor-associated marker TP63 and the proliferation marker Ki67 (Fig. 4b). These results indicate that NGFR expression is more closely associated with an undifferentiated, proliferative cell state than with a differentiated state.

**Figure 4.**
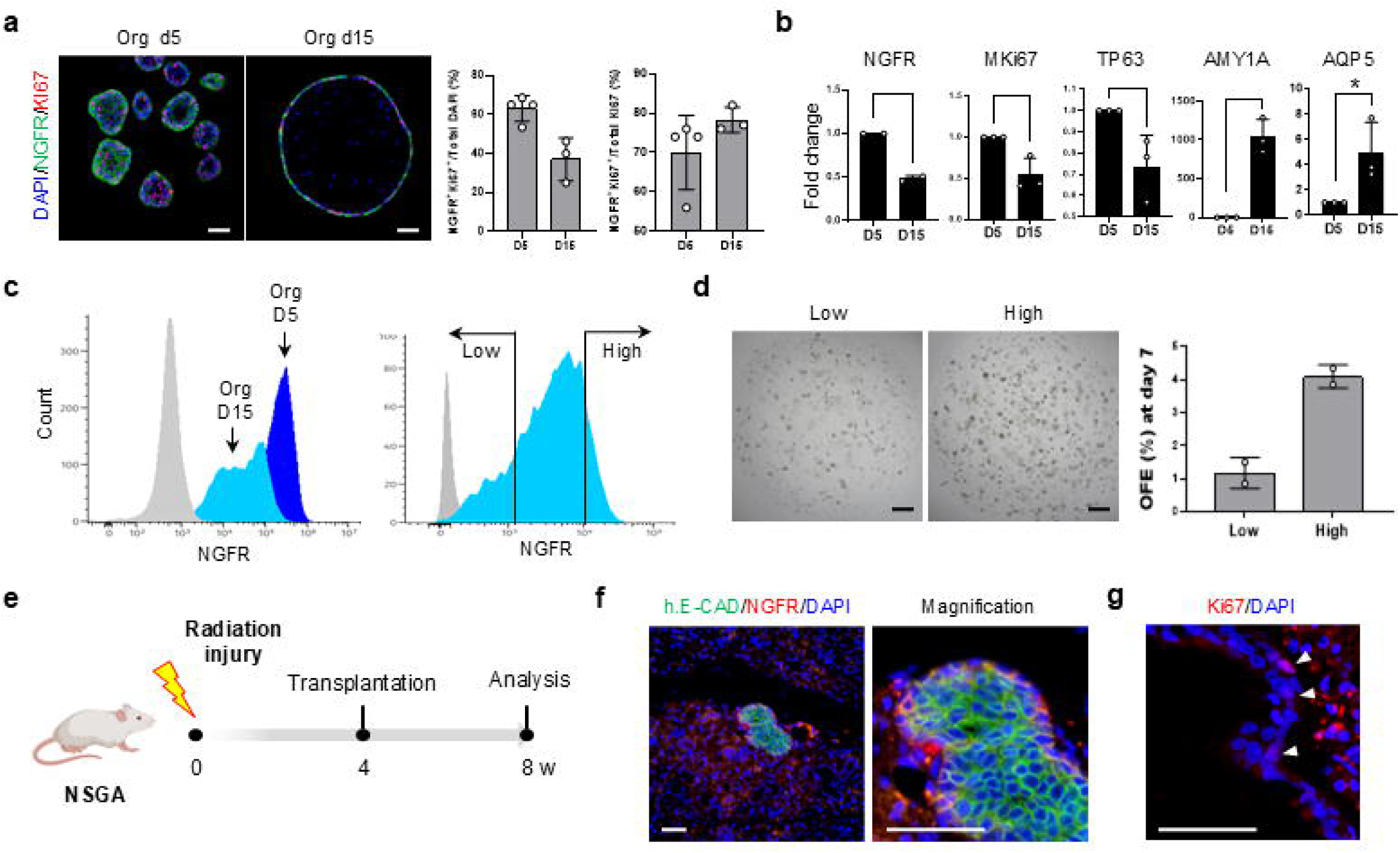
NGFR expression marks a progenitor-enriched organoid state with engraftment potential. a. Immunofluorescence images of human salivary gland organoids stained for NGFR, Ki67, and DAPI at day 5 and day 15. Quantification shows the proportion of NGFR^+^Ki67^+^ cells among total DAPI^+^ cells and among total Ki67^+^ cells at the indicated time points. Scale bar, 50 μm. b. qRT-PCR analysis of NGFR, the proliferation marker MKi67, the basal/progenitor-associated marker TP63, and the acinar differentiation markers AMY1A and AQP5 in human salivary gland organoids at day 5 and day 15. Expression is shown as fold change relative to day 5. *p < 0.05. c. Flow cytometry histograms of NGFR expression in salivary gland organoids harvested at day 5 and day 15 (left). Based on the distribution of NGFR expression at day 15, cells were gated into NGFR^low^ and NGFR^high^ fractions for fluorescence-activated cell sorting (right). d. Bright-field images and quantification of organoid-forming efficiency (OFE) at day 7 after reseeding sorted NGFR^low^ and NGFR^high^ cells from day 15 organoids. Data are shown as mean ± SD. Scale bar, 500 μm. e. Experimental scheme for radiation-induced injury and transplantation of human salivary gland organoids in NSGA mice, with analysis at the indicated time points. f. Immunofluorescence staining of engrafted human salivary gland organoids at 4 weeks after transplantation, stained for human E-cadherin, NGFR, and DAPI. A magnified view of the engrafted organoid structure is shown on the right. Scale bars, 50 μm. g. Ki67 immunostaining of engrafted organoid structures, showing proliferating cells along the outer epithelial layer. Scale bar, 50 μm.

We then tested whether NGFR could serve as a practical surface marker for cell isolation in organoid cultures. Day 15 organoids were dissociated into single cells and sorted into NGFR^low^ and NGFR^high^ populations based on expression intensity, followed by secondary organoid culture under identical conditions (Fig. 4c). The NGFR^high^ fraction showed significantly higher secondary organoid-forming efficiency than the NGFR^low^ fraction (Fig. 4d). This finding suggests that NGFR expression level is functionally associated with a progenitor-enriched, organoid-forming state in salivary gland organoids.

Finally, to assess the engraftment and structural organization potential of NGFR-enriched organoids, we transplanted human salivary gland organoid-derived cells into a radiation-induced salivary gland injury model in immunodeficient mice (Fig. 4e). Four weeks after transplantation, engraftment was confirmed by human E-cadherin expression, and NGFR expression was observed in the outer layer of the engrafted organoids. Notably, this arrangement resembled the basal ductal organization of native salivary gland ducts (Fig. 4f). In addition, a subset of NGFR^+^ cells expressed Ki67, suggesting that they retained proliferative activity after engraftment (Fig. 4g). Collectively, these results show that NGFR marks a progenitor-enriched organoid state with enhanced secondary organoid-forming capacity and that NGFR-enriched organoids can engraft and organize into duct-like epithelial structures in injured salivary gland tissue.

### Ngfr marks an injury-responsive epithelial population enriched for organoid-forming activity in mouse salivary glands

We next sought to investigate the behavior of Ngfr^+^ cells under in vivo injury conditions. To this end, we first validated Ngfr expression in the mouse salivary gland using approaches analogous to those used for human samples. In the mouse salivary gland, Ngfr expression showed limited overlap with acinar and myoepithelial compartments, but was detected in the ductal compartment. We further examined Ngfr expression in mouse salivary gland organoids and confirmed that Ngfr^+^ cells were maintained during organoid culture (Fig. 5a). To assess the functional properties of Ngfr^+^ cells, we dissociated mouse salivary gland tissue into single cells and performed an organoid-forming assay. Ngfr^+^ cells represented approximately 3% of dissociated mouse salivary gland cells (Fig. 5b). When Ngfr^+^ and Ngfr^-^ fractions were isolated and cultured under identical organoid-forming conditions, organoids were generated primarily from the Ngfr^+^ fraction, with an organoid-forming efficiency of approximately 5%, whereas organoid formation was rarely observed in the Ngfr^-^ fraction (Fig. 5c). These results indicate that organoid-forming activity is enriched within the Ngfr^+^ fraction, rather than being broadly distributed across Ngfr^-^ cells.

**Figure 5.**
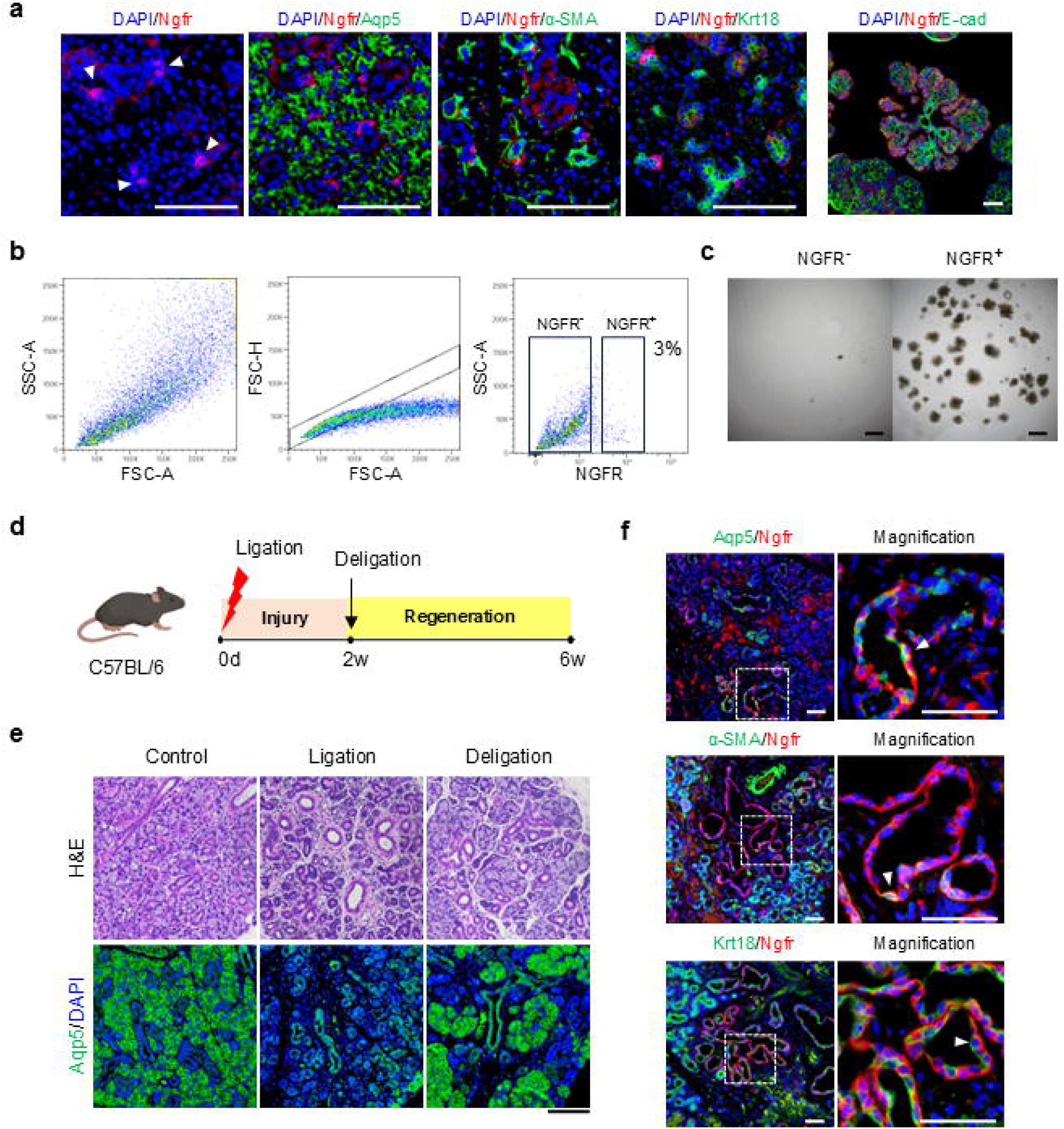
Ngfr marks duct-associated epithelial cells during mouse salivary gland injury and regeneration. a. Immunofluorescence images of mouse salivary gland tissue stained for Ngfr alone or together with Aqp5, α-SMA, or Krt18, and of mouse salivary gland organoids stained for Ngfr and E-cadherin. Arrowheads indicate Ngfr^+^ cells. Scale bars, 50 μm. b. Flow cytometric gating strategy for isolating Ngfr^−^ and Ngfr^+^ cells from dissociated mouse salivary gland tissue. Ngfr^+^ cells represented approximately 3% of the dissociated cell population. c. Bright-field images of organoids derived from sorted Ngfr^-^ and Ngfr^+^ cells. Scale bar, 500 μm. d. Schematic of the duct ligation/deligation injury model used to assess salivary gland injury and regeneration. e. H&E staining of salivary glands from control, ligation, and deligation groups (top), and corresponding immunofluorescence staining for Aqp5 (bottom). Scale bar, 100 μm. f. Immunofluorescence images of salivary glands after duct ligation stained for Ngfr with Aqp5, α-SMA, or Krt18. Higher-magnification views of the boxed regions are shown on the right. Arrowheads indicate cells showing co-expression of Ngfr with the indicated markers. Scale bar, 50 μm.

We then applied a duct ligation model to determine how Ngfr^+^ cells respond to injury. After identifying the main salivary duct by KRT5 expression, we induced salivary gland injury by duct ligation for approximately 2 weeks (Fig. 5d, Supplementary Fig. 3a). Following ligation, we observed marked acinar cell loss and morphological changes in the ductal compartment (Fig. 5e, Supplementary Fig. 3b). Although the overall ductal structure was largely preserved, ducts appeared more condensed within the same tissue area. Upon deligation, acinar cells increased during the regeneration phase, indicating partial recovery of the acinar compartment (Fig. 5e). Notably, strong Ngfr expression emerged in a subset of the remaining ductal structures after ligation (Fig. 5f). Some Ngfr^+^ cells were observed in close proximity to, or partially overlapping with myoepithelial or acinar cell markers. These observations suggest that, following injury, Ngfr^+^ cells are associated with remodeling ductal regions and an injury-associated epithelial state.

To determine whether Ngfr responsiveness was restricted to duct ligation, we next applied an additional local inflammatory injury condition by directly administering Concanavalin A into the salivary gland (Supplementary Fig. 3c). Two weeks after Concanavalin A treatment, localized tissue damage was observed within the salivary gland, accompanied by a focal reduction in mucin staining and the acinar marker Aqp5 (Supplementary Fig. 3d, e). In contrast, Ngfr expression was locally increased around the injured region, and a subset of Ngfr^+^ cells was associated with the proliferation marker Ki67 (Supplementary Fig. 3f). Together, these findings indicate that Ngfr marks a rare epithelial population whose isolated Ngfr^+^ fraction is enriched for organoid-forming activity and that Ngfr expression is associated with injury-responsive ductal regions after salivary gland damage.

### Ngfr-lineage cells contribute to ductal and acinar epithelial compartments during post-injury regeneration

We next sought to determine whether Ngfr-expressing cells contribute to epithelial compartments during regeneration after injury. To this end, we generated an inducible Ngfr-CreERT2 knock-in mouse using a CRISPR-Cas9-based strategy (Fig. 6a). Ngfr-CreERT2 mice were crossed with Rosa26-tdTomato reporter mice to generate double-heterozygous mice for lineage tracing (Fig. 6b). Following tamoxifen administration to label Ngfr-expressing cells, salivary gland injury was induced by duct ligation. Two weeks later, the ligation clip was removed to allow regeneration, and tissues were analyzed after an additional 4-week recovery period to assess the contribution of Ngfr-lineage cells (Fig. 6c).

**Figure 6.**
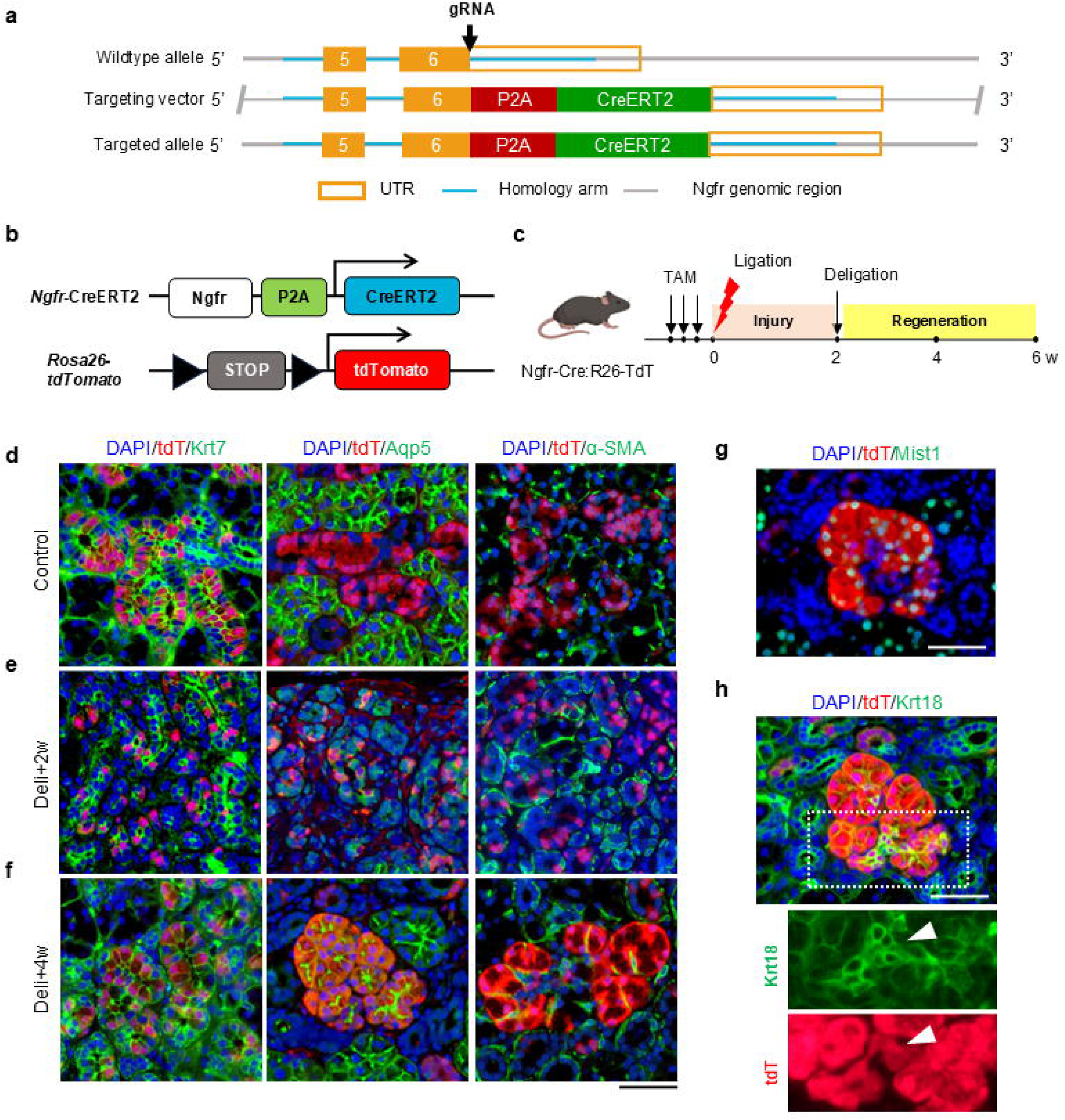
Ngfr-lineage tracing reveals ductal and acinar epithelial contribution after salivary gland injury. a. Schematic of the Ngfr-CreERT2 knock-in strategy showing targeted insertion of P2A-CreERT2 into the endogenous Ngfr locus. b. Genetic strategy for lineage tracing using Ngfr-CreERT2; Rosa26-tdTomato reporter mice. c. Experimental scheme for tamoxifen-inducible lineage tracing during duct ligation-induced salivary gland injury and post-deligation regeneration. Tamoxifen was administered before duct ligation to label Ngfr-expressing cells and their descendants. d, e, f Representative immunofluorescence images of salivary glands from Ngfr-CreERT2; Rosa26-tdTomato mice under control conditions (d), 2 weeks after deligation (e), and 4 weeks after deligation (f). Sections were stained for tdTomato with Krt7, Aqp5, or α-SMA, with DAPI counterstaining. tdTomato marks Ngfr-lineage cells. Scale bar, 50 μm. g. Representative immunofluorescence image of regenerating salivary gland tissue 4 weeks after deligation stained for tdTomato, Mist1, and DAPI. Scale bar, 100 μm. h. Representative immunofluorescence image of regenerating salivary gland tissue 4 weeks after deligation stained for tdTomato, Krt18, and DAPI. Channel-separated higher-magnification views of the boxed region are shown below. Scale bar, 100 μm.

Under homeostatic conditions, tdTomato^+^ cells were observed predominantly within the KRT7^+^ ductal compartment, suggesting that the Ngfr lineage is maintained mainly within the ductal epithelial cells (Fig. 6d). At 2 weeks after deligation following duct ligation injury, tdTomato^+^ cells remained predominantly associated with the KRT7^+^ ductal compartment, with only limited co-expression with the acinar marker AQP5 (Fig. 6e). By 4 weeks after deligation, tdTomato^+^ cells were detected in both the KRT7^+^ ductal and AQP5^+^ acinar compartments, indicating increased acinar contribution of Ngfr-lineage cells during regeneration (Fig. 6f). Co-localization with the myoepithelial marker α-SMA was not observed under these conditions. A subset of tdTomato^+^ cells in the regenerating acinar region also expressed MIST1, a marker associated with mature acinar differentiation (Fig. 6g). In addition, KRT18 and tdTomato staining revealed that, in some regions, tdTomato^+^ cells were distributed along continuous epithelial structures extending from ductal regions toward acinar regions (Fig. 6h). This pattern suggests that Ngfr-lineage cells may participate in epithelial regeneration associated with duct–acinar compartment reorganization.

We next asked whether Ngfr-expressing cells in other organs could contribute to epithelial compartments, as observed in the salivary gland. We first surveyed multiple glandular organs, including the parotid gland, pancreas, mammary gland, lacrimal gland, and uterus, and found Ngfr expression within or adjacent to E-cadherin^+^ epithelial compartments (Supplementary Fig. 4a). We therefore performed tamoxifen-inducible lineage tracing using Ngfr-CreERT2; Rosa26-tdTomato mice to determine whether Ngfr-expressing cells and their descendants contribute to epithelial compartments in vivo (Supplementary Fig. 4b). Six weeks after tamoxifen induction, tdTomato^+^ Ngfr-lineage cells were broadly detected in the uterine epithelium and were also observed in selected mucosal and surface epithelial tissues, including the tongue, eye, and trachea (Supplementary Fig. 4c). These observations suggest that Ngfr-lineage contribution to epithelial compartments is not restricted to the salivary gland and can be observed in selected glandular, mucosal, and surface epithelial tissues under homeostatic conditions.

Collectively, these findings indicate that Ngfr-lineage cells are largely maintained within the ductal compartment under homeostatic conditions but contribute to both ductal and acinar epithelial compartments during post-injury regeneration. Ngfr-lineage contribution was also observed in selected glandular, mucosal, and surface epithelial tissues under homeostatic conditions.

## Discussion

Severe salivary gland injury often results in persistent dysfunction with limited endogenous recovery. Identifying epithelial populations that participate in repair is critical for understanding salivary gland regeneration and developing cell-based repair strategies, highlighting the need for markers that enable precise identification and isolation of these cells. In this study, we identify NGFR as a surface marker of a restricted basal duct-associated epithelial population in the human salivary gland and enriches fractions with organoid-forming activity. In parallel, mouse injury models and lineage tracing show that Ngfr marks an injury-responsive epithelial population whose lineage contributes to ductal and acinar epithelial compartments during post-injury regeneration. By integrating human single-cell transcriptomics, prospective cell isolation, organoid-based functional assays, and mouse injury lineage tracing, this study links NGFR/Ngfr-marked basal duct-associated epithelial identity to both ex vivo organoid-forming activity and in vivo epithelial contribution after injury.

Previous markers have provided an important foundation for studying salivary gland progenitors, but limitations remain with respect to isolation, validation in human tissue, and cellular specificity. In this study, we used scRNA-seq analysis to identify a new surface marker that is distinct from conventional basal markers. KRT5/CK5 has been widely used as a representative marker of basal ductal cells in the salivary gland. However, KRT5 is also broadly observed in the myoepithelial compartment, limiting its utility for isolation of a specific basal duct subpopulation (May et al., 2018). In contrast, NGFR showed limited overlap with major differentiated cell markers and was relatively enriched in a basal duct-associated subpopulation. Thus, NGFR may represent a more suitable candidate surface marker for prospectively isolating a basal duct-associated epithelial population from human salivary gland tissue.

Beyond its spatial specificity, the functional relevance of NGFR was supported by organoid-forming assays. Because organoid formation provides a functional readout of epithelial growth capacity under defined culture conditions, it allowed us to compare the activity of prospectively isolated cell fractions. Although NGFR^+^ cells represented a minor fraction of human salivary gland tissue, organoid-forming activity was markedly enriched within the NGFR^+^ fraction compared with the NGFR^-^ fraction. In addition, NGFR^+^EPCAM^+^ cells showed higher organoid-forming efficiency than CD24^+^EPCAM^+^ and ITGA6^+^EPCAM^+^ fractions under the same culture conditions. These findings indicate that NGFR can be used to enrich human salivary epithelial fractions with enhanced organoid-forming capacity. However, these data should be interpreted as enrichment of organoid-forming activity within the NGFR^+^ fraction, rather than evidence that all NGFR^+^ cells are organoid-initiating stem cells. Future single-cell-derived organoid assays, limiting dilution analysis, and serial passaging experiments will be important to define the frequency and self-renewal properties of NGFR-enriched organoid-forming cells.

During organoid culture, NGFR expression was retained in highly proliferative cells and declined with differentiation. These observations suggest that NGFR^+^ basal duct cells do not represent a constitutively proliferative population, but rather a progenitor-like epithelial state that can be activated or maintained under culture conditions. Consistent with this interpretation, NGFR^high^ cells isolated from organoids showed enhanced secondary organoid-forming capacity compared with NGFR^low^ cells. These findings support the idea that NGFR expression marks a progenitor-enriched organoid state associated with renewed organoid-forming activity. Together, the tissue-sorting and organoid-resorting experiments suggest that NGFR is useful not only for prospective isolation from primary human salivary gland tissue, but also for tracking functionally enriched epithelial states during organoid expansion. Defining the niche signals that regulate this state, including WNT, EGF/FGF, Notch, YAP/TAZ, and neurotrophin-related pathways, may provide important insight into the mechanisms that control epithelial plasticity and regenerative failure in the injured salivary gland.

NGFR-enriched organoids engrafted in injured salivary gland tissue and formed duct-like epithelial structures with NGFR^+^ cells positioned along the outer layer. This arrangement resembled the basal ductal organization of native salivary gland ducts, indicating that NGFR^+^ cells retain the capacity to establish duct-like structure in an injured tissue environment. Importantly, the current transplantation data demonstrate engraftment and duct-like organization, but do not establish broad acinar reconstruction or functional restoration of salivary secretion. Thus, full epithelial reconstruction may require not only the intrinsic growth and organization capacity of NGFR^+^ cells, but also efficient host niche integration and appropriate acinar differentiation cues. Future studies should optimize delivery methods and niche-supporting strategies to improve engraftment, acinar maturation, and functional tissue regeneration.

Together with the human data, the mouse injury models extend the relevance of NGFR/Ngfr-marked epithelial states from organoid-forming activity to tissue remodeling after salivary gland injury. This provided the rationale for using Ngfr-CreERT2 lineage tracing to determine whether Ngfr-expressing cells contribute to epithelial regeneration *in vivo*. Ngfr-CreERT2 lineage tracing demonstrated that Ngfr-lineage cells contribute to both ductal and acinar epithelial compartments during regeneration after injury. In some regions, Ngfr-lineage cells were distributed along continuous epithelial structures spanning ductal and acinar regions, suggesting that the Ngfr lineage may participate in duct–acinar compartment reorganization rather than appearing only as scattered lineage-labeled cells. These data support a model in which Ngfr-expressing duct-associated epithelial cells acquire injury-responsive behavior and contribute to epithelial repair after damage. However, these data do not define the direct differentiation route of Ngfr-lineage cells or establish clonal duct–acinar unit formation. Clonal lineage tracing, three-dimensional reconstruction, and time-course analysis will be required to further define the cellular dynamics of Ngfr-lineage contribution. Beyond the salivary gland, Ngfr-lineage cells were also detected in epithelial compartments of selected glandular, mucosal, and surface epithelial tissues under homeostatic conditions. Although this observation suggests that Ngfr-lineage epithelial contribution may not be restricted to the salivary gland, tissue-specific functional studies will be required to determine whether these cells support epithelial regeneration after injury.

A key strength of this study is that NGFR/Ngfr connects multiple layers of evidence across human tissue, organoid culture, transplantation, injury response, and lineage tracing: spatial restriction within a basal duct-associated compartment, functional enrichment in organoid-forming assays, engraftment-associated duct-like organization, injury responsiveness, and lineage contribution to ductal and acinar epithelial compartments. The ability to identify and isolate this epithelial population provides an entry point for defining the signaling and niche cues that regulate injury-responsive epithelial plasticity, including the potential role of NGFR/Ngfr-associated neurotrophin pathways in salivary gland repair. Future studies that modulate this marker-defined epithelial state may help develop strategies to enhance epithelial regeneration after salivary gland injury.

In summary, this study identifies NGFR/Ngfr as a conserved marker of an injury-responsive basal duct-associated epithelial population. By linking prospective isolation, organoid-forming activity, and ductal–acinar epithelial contribution after injury, our findings establish a cellular basis for harnessing injury-responsive epithelial states to promote salivary gland repair and regeneration.

## Methods

### Human salivary gland samples

This study was approved by the Institutional Review Board of Konkuk University Hospital (IRB No. 202011062, 2024-02-009-001). Histologically normal salivary gland tissues were collected from normal regions of patients with head and neck cancer.

### Mice

All animal experiments were approved by the Red-Green Bioconvergence Institute Institutional Animal Care and Use Committee (IACUC approval no. IACUC-RGB-260005) and were performed in accordance with institutional guidelines for animal care and use. Ngfr-CreERT2 mice were crossed with Rosa26-tdTomato reporter mice and used for lineage tracing experiments. Eight-week-old female C57BL/6J mice were used for organoid generation and injury modeling. Eight-week-old female NOD-Prkdc^em1Baek^Il2rg^em1Baek^ (NSGA) mice were used for transplantation experiments. All mice were housed in a constant temperature (approximately 23 °C) room on a 12-h light–dark cycle and fed *ad libitum*. Body weight was measured every 2 weeks throughout the experiments. All surgical procedures on mice were conducted under anesthesia using a combination of 100 mg/kg ketamine and 10 mg/kg xylazine administered intraperitoneally or using isoflurane inhalation. Mice were euthanized by carbon dioxide (CO2) inhalation.

### Generation of Ngfr-CreERT2 knock-in mice

The Ngfr (C terminal-CreERT2) knock-in mouse was generated by Cyagen Company (Suzhou, China). Using the BAC clone RP23-67E18 as a template, homology arms were amplified by PCR. The mouse Ngfr gene (NCBI Reference Sequence: NM_033217.3) is located on mouse chromosome 11. Six exons have been identified, with the ATG start codon in exon 1 and the TGA stop codon in exon 6. For the KI model, the TGA stop codon was replaced with “P2A-CreERT2” cassette. The homology arms and the C terminal-CreERT2 were constructed into conventional mammalian expression vectors using In-Fusion cloning technology. Cas9 protein, two gRNAs, and a donor vector were co-injected into fertilized eggs. The embryos were transferred to a recipient female mouse to obtain F0 mice. The genotype of the Ngfr (C terminal-CreERT2) knock-in mice was confirmed by PCR using specific primers and sequencing.

### Salivary gland cell isolation and fluorescence-activated cell sorting

Human and mouse salivary gland cells were isolated and cultured as previously described (Jeon et al., 2024). Briefly, tissue samples were weighed to adjust the digestion enzyme volume, rinsed three times with PBS, and mechanically minced into small fragments. The minced tissues were then enzymatically dissociated in HBSS/1% BSA buffer containing 0.63 mg/mL collagenase II (Invitrogen) and 0.5 mg/mL hyaluronidase (Sigma-Aldrich) for 2 h at 37 °C with shaking at 120 rpm. Dissociated human salivary gland single cells were resuspended in DPBS containing 2% FBS and filtered through a strainer. Cells were incubated with Human BD Fc Block (BD Biosciences, 564219) or Mouse BD Fc Block™ (BD Biosciences, 553141) for 15 min at room temperature and stained for 30 min at 4°C with EPCAM-FITC together with NGFR-APC, CD24-APC, or ITGA6-APC in separate staining reactions. After washing, cells were resuspended in culture medium and sorted using a BD FACSMelody™ Cell Sorter (BD Biosciences). Debris and doublets were excluded, and singlet cells were gated for downstream sorting. Cells were sorted as NGFR^+^ and NGFR^−^ fractions or as EPCAM^+^NGFR^+^, EPCAM^+^CD24^+^, and EPCAM^+^ITGA6^+^ fractions depending on the experiment. Sorted cells were collected in culture medium supplemented with 10 μM Y-27632 and seeded for organoid-forming assays at 1,000 to 3,000 cells per well. For sorting of human salivary gland organoid-derived cells, organoids were dissociated into single cells using 0.25% trypsin-EDTA and processed using the same staining and sorting procedure. After exclusion of debris and doublets, cells were sorted into NGFR^high^ and NGFR^low^ fractions. Sorted organoid-derived cells were collected in culture medium supplemented with 10 μM Y-27632 and seeded at 2,000 to 3,000 cells per well for secondary organoid-forming assays. Flow cytometry data were analyzed using BD FACSChorus™ software and FlowJo™ v11.1.1 software (BD Life Sciences).

### Salivary gland organoid culture

Organoid culture was performed as previously described with minor modifications (Jeon et al., 2024). Briefly, 5,000 dissociated salivary gland cells were suspended in a 20-μL mixture of Matrigel and culture medium (1:1, v/v), plated as domes in 48-well plates, and allowed to polymerize at 37°C in a 5% CO2 incubator. Human salivary gland organoids were cultured in expansion medium (EM); AdDMEM/F12 (GIBCO) supplemented with Glutamax, HEPES, 100 U/mL Penicillin-Streptomycin (all Thermo Fisher scientific), 1.25 mM N-acetylcysteine (Sigma-Aldrich) and the following growth factors: 100 ng/mL Noggin, 10% R-spondin 1 conditioned medium (Trevigen), 0.5 μM A83-01 (Tocris), 5 nM neuregulin, 10 μM Y27632 and 100 ng/mL FGF10 (Peprotech). Mouse salivary gland organoids were cultured in DMEM/F12 (GIBCO) containing Pen/Strep antibiotics (Invitrogen), Glutamax (Invitrogen), 20 ng/mL epidermal growth factor (EGF; Sigma-Aldrich), 20 ng/mL fibroblast growth factor-2 (FGF2; Sigma-Aldrich), N2 (Invitrogen), 10 μg/mL insulin (Sigma-Aldrich), 1 μM dexamethasone, 50% Wnt3A, R-spondin and noggin conditioned medium (ATCC). For passage, organoids at day 5 were collected and incubated at 37 °C for 5 min with 0.25% trypsin-EDTA and dissociated into single cells.

### Histological analysis

Tissues and organoids were fixed overnight in 4% formaldehyde at 4°C, processed, and embedded in paraffin. Paraffin sections were cut at a thickness of 4 μm, deparaffinized in xylene, and rehydrated through a graded ethanol series. Hematoxylin and eosin staining was performed using a standard protocol with a commercial staining kit (Abcam, cat# ab245880). For mucin detection, Alcian blue staining was performed using an Alcian blue staining kit (Agilent, cat# AR16092). Briefly, sections were incubated in acetic acid solution for 3 min at room temperature, followed by staining with Alcian Blue solution (pH 2.5) for 30 min. After washing, sections were counterstained with Nuclear Fast Red solution for 5 min. Images were acquired using a TS2-LS microscope (Nikon).

### Immunofluorescence analysis

Paraffin-embedded tissue and organoid sections were deparaffinized in xylene and rehydrated through a graded ethanol series. Antigen retrieval was performed in sodium citrate buffer (10 mM sodium citrate, 0.05% Tween 20, pH 6.0) at 95°C for 30 min. Sections were blocked with PBS containing 5% normal horse serum (Vector Laboratories) for 1 h at room temperature and then incubated overnight at 4°C with the primary antibodies listed in the table. After washing, sections were incubated with secondary antibodies at a 1:500 dilution for 1 h at room temperature. Nuclei were counterstained with Hoechst 33342 (1 μg/mL; Sigma), and fluorescence images were acquired using a TS2-LS microscope (Nikon).

### CODEX antibody conjugation

Antibody barcode conjugation was performed using the antibody conjugation kit (Akoya Biosciences, #7000009) according to the manufacturer’s instructions. Selected antibodies were conjugated to their corresponding CODEX barcodes. Antibody–barcode pairings were AQP5-BX020, CK19-BX035, NGFR-BX006, CK5-BX010, CK7-BX016, TUJ1-BX034, LCN2-BX040, α-amylase-BX030, MUC5B-BX049, and LPO-BX002. Briefly, CODEX barcodes were resuspended in molecular biology-grade water, mixed with Conjugation Solution, and incubated with the corresponding antibodies for 2 h at room temperature. Barcode-conjugated antibodies were purified using Purification Solution, recovered in Antibody Storage Solution, and stored at 4°C before staining. Conjugated antibodies were validated by gel electrophoresis according to the Akoya Biosciences guidelines before use.

### Multiplex staining

Formalin-fixed paraffin-embedded tissue sections were baked at 65°C for 1 h, deparaffinized twice in xylene, and rehydrated through a graded ethanol series of 100%, 100%, 90%, 80%, 70%, 50%, and 30% ethanol, followed by two washes in ddH2O. Slides were fixed with 4% paraformaldehyde solution in PBS (T&I, BPP-9004) for 1 h, washed with ddH2O, and subjected to antigen retrieval using 1× AR9 buffer prepared from 10× AR9 Solution (Akoya Biosciences, #AR9001KT). Antigen retrieval was performed in a pressure cooker at high pressure (∼11.6 PSI, 110°C) for 20 min, followed by cooling at room temperature for 1 h. After antigen retrieval, slides were washed with ddH2O and equilibrated sequentially in Hydration Buffer and Staining Buffer (Akoya Biosciences, #7000017). The antibody cocktail was prepared using the 41 antibodies listed in the table. Briefly, the antibody cocktail solution was prepared with Staining Buffer, N-Blocker, G-Blocker, J-Blocker, and S-Blocker (Akoya Biosciences, #7000017), and the appropriate volume of each PhenoCycler antibody was added to generate the final staining solution. The prepared antibody cocktail staining solution was applied to the tissue section to fully cover the sample, and slides were incubated for 3 h at room temperature on a rocker set to 30 rpm. After staining, slides were post-fixed with 1.6% PFA Post-Staining Fixing Solution and PhenoCycler Fixative Reagent (Akoya Biosciences, #7000017), washed with 1× PBS, and processed for PhenoCycler-Fusion imaging.

### Multiplex staining imaging

Reporter Stock Solution for 16 imaging cycles was prepared in a 15-mL amber tube using 1× Buffer for PhenoCycler with Buffer Additive (Akoya Biosciences, #700019), Assay Reagent (Akoya Biosciences, #7000002), and Nuclear Stain (Akoya Biosciences, #7000003), to a final volume of 4.8 mL. Reporter Master Mixes were prepared for each cycle by adding 5 μL of the assigned antibody reporters to the Reporter Stock Solution, with a final volume of 250 μL per cycle. Each Reporter Master Mix was transferred to the corresponding well of a 96-well plate (Akoya Biosciences, #7000006), sealed with a foil plate seal (Akoya Biosciences, #7000007), and stored protected from light until use. The experimental plan was set up in PhenoCycler Designer and Fusion software, and all required PhenoCycler reagents and buffers were prepared according to the manufacturer’s instructions.

### Quantitative real-time PCR

Human salivary gland organoids were harvested on days 5 and 15. Total RNA was extracted using TRIzol reagent (Invitrogen) according to the manufacturer’s protocol. For each sample, 1 μg of total RNA was reverse-transcribed into cDNA using PrimeScript RT Master Mix (TAKARA). Quantitative real-time PCR was performed with TB Green Premix (Takara Bio) using a Thermal Cycler Dice Real-Time System III (Takara Bio). Relative mRNA expression was determined using the comparative Ct method.

### Radiation-induced salivary gland injury and organoid transplantation

NSGA mice were subjected to local salivary gland irradiation using a medical linear accelerator (Clinac iX) delivering 6-MV photons at a dose rate of 6 Gy/min. Under anesthesia, the salivary gland region was locally irradiated with a single 7.5-Gy dose of X-rays. Four weeks after irradiation, human salivary gland organoids were dissociated into single-cell suspensions for transplantation. Recipient mice were anesthetized, and the submandibular glands were exposed through a 5-mm anterior cervical incision. Dissociated human salivary gland organoid-derived cells (1 × 10^5^ cells in 20 μL) were injected locally into both glands using a 29-gauge Hamilton syringe. After injection, the incision was closed with sutures.

### Concanavalin A-induced local inflammatory injury

To induce local inflammatory injury, 8-week-old adult C57BL/6J mice were anesthetized, and the salivary glands were exposed through a cervical incision. 40 mg/mL Concanavalin A (ConA; Sigma, C5275) was injected directly into multiple sites within the salivary gland parenchyma. Two weeks after injection, salivary glands were harvested and fixed in 4% paraformaldehyde for further analysis.

### Salivary duct ligation and lineage tracing

For lineage tracing experiments, eight-week-old female Ngfr-CreERT2; Rosa26-tdTomato mice were used. Before duct ligation, mice received tamoxifen (Sigma, T5648) dissolved in corn oil by intraperitoneal injection once daily for five consecutive days at a dose of 75 mg/kg in a volume of 100 μL. For salivary gland duct ligation, mice were anesthetized by isoflurane inhalation. After a cervical incision, the salivary gland was exposed, and the main duct of the left salivary gland was ligated using a titanium hemostatic clip (J9180, VITALITEC) and hemostatic clip appliers (PJ120, VITALITEC). Two weeks after ligation, the clip was carefully removed without causing additional damage to the duct.

### scRNA-seq experiments

Two normal human submandibular gland tissue samples were used for scRNA-seq analysis, designated as Sample 1 and Sample 2, respectively. They were resuspended in PBS with 1% bovine serum albumin. Single-cell RNA-sequencing libraries were then prepared using the Chromium Next GEM Single Cell 3’ reagent kit v3.1 (10X Genomics, Pleasanton, CA) in accordance with the manufacturer’s protocol. Briefly, the cells were diluted into the Chromium Next GEM Chip G to yield a recovery of ∼ 5,000 single cells. Following the library preparation, the libraries were sequenced in multiplex on the Novaseq 6000 sequencer (Illumina, San Diego, CA) to produce on average a minimum of 30,000 reads per single cell.

### Analysis of scRNA-seq data

Sequencing reads were processed with the Cell Ranger version 3.0.1 (10X Genomics, CA) using the Human reference transcriptome GRCh38 from Ensembl. From the gene expression matrix, the downstream analysis was carried out with R version 3.6.2. Quality control, filtering, data clustering and visualization, and the differential expression analysis was carried out using WinSeurat program (Ebiogen Inc., Seoul, Korea) based on Seurat version 3.1.2 R package (Butler et al., 2018) with some custom modifications to the standard pipeline. Additional quality control filtering was applied to remove low-quality cells based on the following criteria: mitochondrial gene ratio > 25%, detected genes < 200 or > 6,000, and total UMI count < 500 or > 50,000, retaining 11,161 high-quality cells for downstream analysis. After scTransform-normalizing the data, the expression of each gene was scaled regressing out the number of UMI and the percent mitochondrial gene expressed in each cell. We performed PCA on the gene expression matrix and used the first 10∼30 principal components for clustering and visualization. Clustering was performed with a resolution of 0.6∼0.8 and visualization was done using UMAP (uniform manifold approximation and projection). Epithelial cells were extracted based on cell type annotation and re-clustered independently using identical parameters (dims = 1:20, resolution = 0.4). Differential gene expression was assessed using FindMarkers (Wilcoxon rank-sum test; adjusted p-value < 0.05). GO Biological Process and KEGG pathway enrichment analyses were performed using clusterProfiler (v4.16.0) with Benjamini-Hochberg correction (adjusted p-value < 0.05). Developmental potency scores were calculated using CytoTRACE2 (v1.0.0) from raw count matrices. Pseudotime trajectory analysis was conducted using Slingshot with the Basal cluster designated as the trajectory origin. Statistical comparisons between NGFR^+^ and NGFR^-^ cells were performed using the Wilcoxon rank-sum test.

### Statistical analysis

Sample sizes and the number of independent experiments are indicated in the figure legends. Data are presented as the mean ± SD unless otherwise indicated. Statistical analyses were performed using GraphPad Prism software version 8.0 (GraphPad Software, CA, USA). For comparisons between two groups, statistical significance was determined using an unpaired two-tailed Student’s t test. For comparisons among multiple groups, one-way analysis of variance (ANOVA) followed by Tukey’s post hoc test was used. For single-cell RNA-seq analyses, differential expression and comparisons between NGFR^+^ and NGFR^−^ cells were assessed using the Wilcoxon rank-sum test, and adjusted p values were calculated using Benjamini-Hochberg correction where applicable. p < 0.05 was considered statistically significant.

## Data availability

Single-cell RNA-seq data generated in this study have been deposited in the Gene Expression Omnibus under accession code GSE330533. The data will be publicly available as of the date of publication. Processed count matrices, cell metadata and cluster annotations are available in the same repository.

## Acknowledgments

This research was supported by a grant of the Korea Health Technology R&D Project through the Korea Health Industry Development Institute (KHIDI), funded by the Ministry of Health & Welfare, Republic of Korea (grant number : HI22C1462). This work was supported by the Technology Innovation Program (or Industrial Strategic Technology Development Program-grant number: 20018578, Production standardization and development of analysis to verify the quality and characterization of organoid based regeneration medicine) funded by the Ministry of Trade, Industry & Energy (MOTIE, Korea). This work was supported by grants from the National Research Foundation of Korea (NRF), funded by the Ministry of Science, ICT, and Future Planning under grant number RS-2026-25515390 and the KRIBB Research Initiative Program under grant number KQM0042611. Biological illustrations in Fig. 3a were created using BioRender.com.

## Author Contributions

S.G.J. and J.Y. designed the project. S.G.J. performed most experiments. D.H.B. performed bioinformatic analyses of the single-cell RNA-sequencing data. S.G.J., J.-Y.P., and T.V.T.N. performed investigation and validation experiments. S.G.J. and M.Y. performed the animal experiments. Y.C.L. provided human salivary gland tissue samples. K.J.L., S.-H.L., and Y.C.L. provided scientific input. S.G.J., D.H.B., and J.Y. wrote the manuscript. E.J.B., M.-Y.S., and J.Y. edited the manuscript. J.Y. supervised the research.

## Figure legends

**Supplementary Figure 1.**
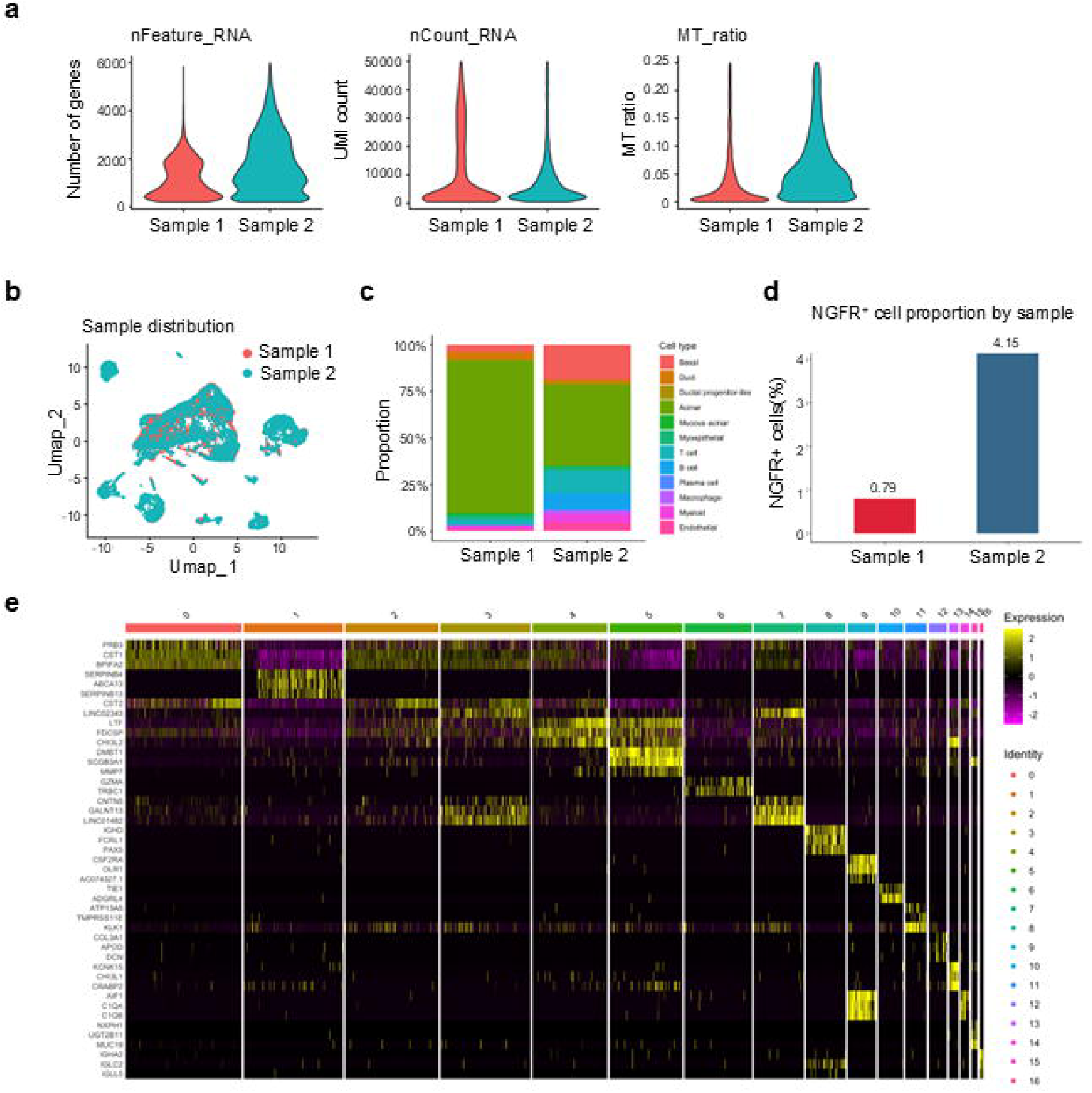
Quality control, sample distribution, and cluster annotation of human salivary gland scRNA-seq data. a. Violin plots showing quality control metrics for cells retained after filtering, including the number of detected genes (nFeature_RNA), total UMI counts (nCount_RNA), and mitochondrial gene ratio (MT_ratio), grouped by sample. **b.** UMAP visualization of 11,161 cells colored by sample origin, showing the distribution of Sample 1 and Sample 2 after integration. **c.** Stacked bar plots showing the proportional distribution of annotated cell types in each sample. **d.** Bar plot showing the proportion of NGFR^+^ cells among total cells in each sample. NGFR^+^ cells were detected in both samples, comprising 0.79% of cells in Sample 1 and 4.15% of cells in Sample 2. **e.** Heatmap showing scaled expression of top differentially expressed marker genes across 17 clusters, supporting the cell type annotation shown in Figure 1.

**Supplementary Figure 2.**
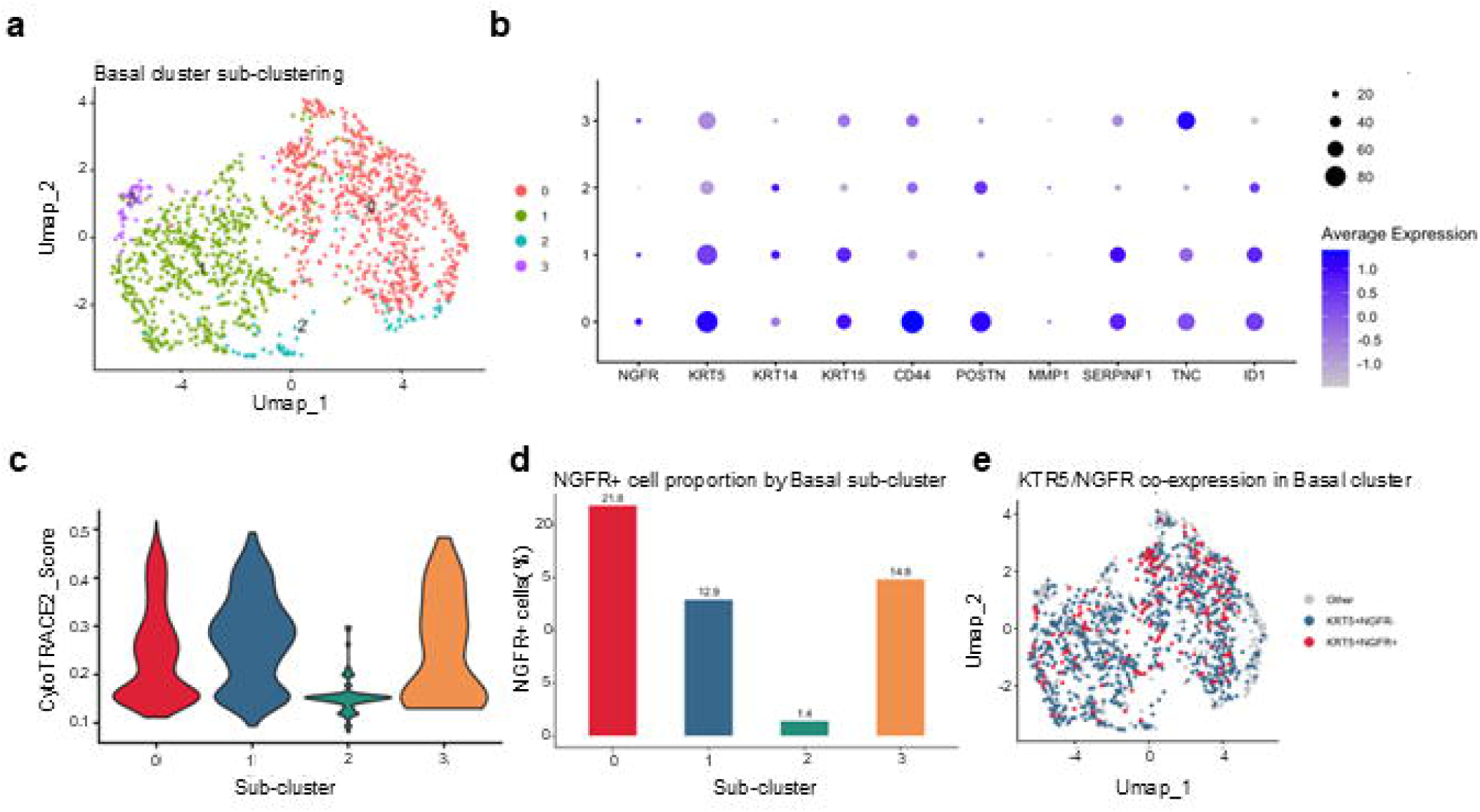
Transcriptional heterogeneity within the basal cluster. a. UMAP visualization of subclustered basal cells, showing four basal subclusters (0-3). b. Dot plot showing expression of selected marker genes across basal subclusters. Dot size indicates the percentage of expressing cells, and color intensity indicates scaled average expression. c. Violin plots comparing CytoTRACE2 developmental potency scores across basal subclusters. Subcluster 3 exhibits the highest potency, while subcluster 2 shows the lowest. d. Bar plot showing the proportion of NGFR^+^ cells within each Basal sub-cluster. Subcluster 0 contains the highest proportion of NGFR^+^ cells (21.8%). e. UMAP visualization of basal cells colored by KRT5 and NGFR co-expression status. KRT5^+^NGFR^+^ cells are shown in red and KRT5^+^NGFR^-^ cells in blue. KRT5^+^NGFR^+^ cells were distributed across multiple basal subclusters rather than being confined to a single subcluster. NGFR expression is detected at low levels consistent with its restricted expression in 16.9% of basal cells as shown in Figure 1E.

**Supplementary Figure 3.**
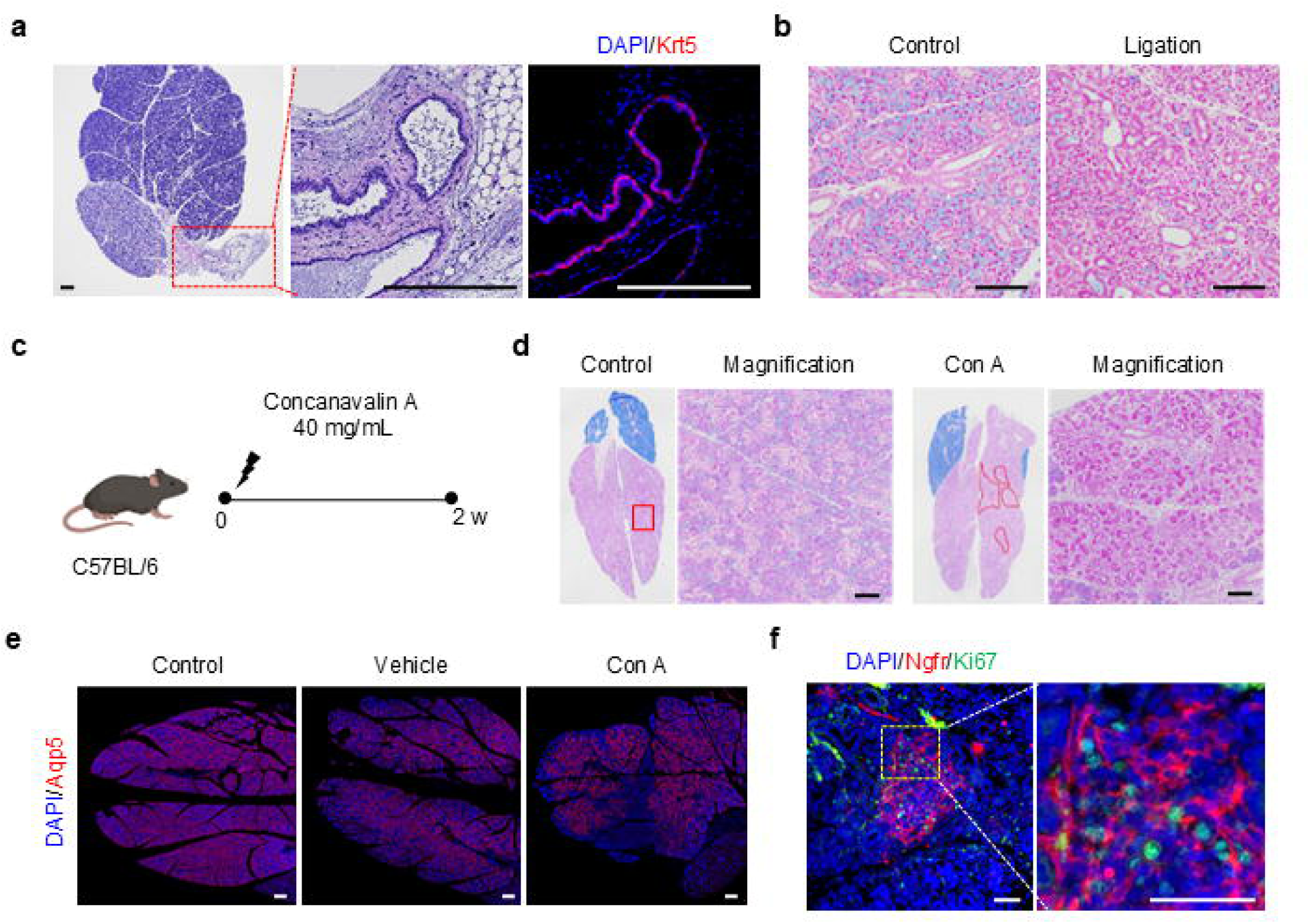
Validation of duct ligation and Concanavalin A-induced injury responses in mouse salivary glands. a. Representative histological and immunofluorescence images of the main salivary duct region used for duct ligation. H&E staining shows the anatomical location of the main duct, and Krt5 immunofluorescence marks the ductal epithelium. Scale bar, 250 μm. b. Alcian blue staining of control and duct-ligated salivary glands 2 weeks after ligation. Mucin-positive regions are reduced after duct ligation. Scale bar, 100 μm. c. Experimental scheme for Concanavalin A-induced local salivary gland injury. Salivary glands were analyzed 2 weeks after Concanavalin A administration. d. Alcian blue staining of control and Concanavalin A-treated salivary glands. Boxed and outlined areas in the low-magnification images are shown at higher magnification in the adjacent panels. Concanavalin A-treated glands show focal tissue damage. Scale bar, 100 μm. e. Whole-gland immunofluorescence staining for Aqp5 in control, vehicle-treated, and Concanavalin A-treated salivary glands. Scale bar, 250 μm. f. Immunofluorescence staining for Ngfr and Ki67 in a focal injury region after Concanavalin A treatment. A higher-magnification view of the boxed region is shown on the right. Scale bar, 50 μm.

**Supplementary Figure 4.**
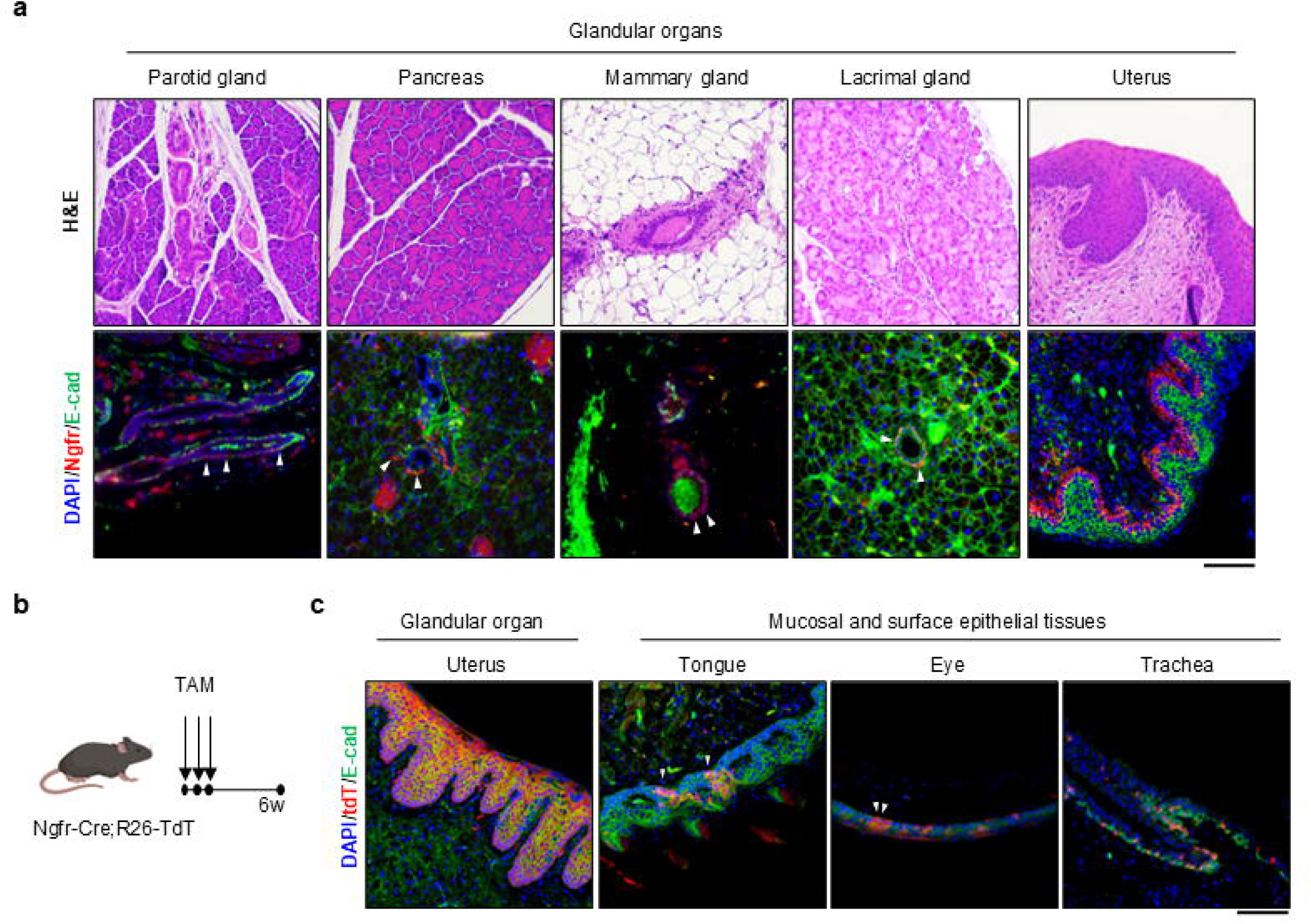
Ngfr-CreERT2-labeled cells are detected in multiple epithelial and glandular organs. a. Representative H&E and immunofluorescence images of Ngfr expression in glandular organs. H&E staining shows tissue morphology in the parotid gland, pancreas, mammary gland, lacrimal gland, and uterus. Immunofluorescence staining for Ngfr and E-cadherin shows the spatial relationship between Ngfr^+^ cells and epithelial compartments in each tissue. Scale bar, 100 μm. b. Experimental design for tamoxifen-inducible lineage tracing in Ngfr-CreERT2; Rosa26-tdTomato mice. b. Representative immunofluorescence staining of the uterus, tongue, eye, and trachea from Ngfr-CreERT2;Rosa26-tdTomato mice 6 weeks after tamoxifen induction. tdTomato marks Ngfr-lineage cells, and E-cadherin marks epithelial compartments. Scale bar, 100 μm.

**Supplementary Table 1.** Antibodies used in this study.

| Antibodies | Company | Cat.number |
| --- | --- | --- |
| rabbit anti-aquaporin 5 | Alomone labs | Cat# AQP-005 |
| mouse anti-cytokeratin 19 | Biolegend | Cat# 628502 |
| rabbit anti-LPO | Sigma Aldrich | Cat# HPA028688 |
| rabbit anti-α-amylase | Cell Signaling | Cat# 3796S |
| rabbit anti-CK5 | Biolegend | Cat# 905504 |
| mouse anti-cytokeratin 7 | Invitrogen | Cat# MA5-11986 |
| mouse anti-TUJ1 | Biolegend | Cat# 801201 |
| goat anti-LCN2 | R&D systems | Cat# AF1757 |
| mouse anti-MUC5B | Abcam | Cat# ab105460 |
| rabbit anti-CD24 | Novus Biologicals | Cat# NBP1-46390SS |
| mouse anti-EPCAM | Cell Signaling | Cat# 2929S |
| rabbit anti-ITGA6 | Abcam | Cat# ab181551 |
| rabbit anti-Ki67 | Abcam | Cat# ab15580 |
| rabbit anti-E-Cadherin | Abcam | Cat# ab40772 |
| mouse anti-NGFR | Santa cruz | Cat# sc-13577 |
| rabbit anti-NGFR | Novus Biologicals | Cat# NBP2-67296 |
| goat anti-NGFR | Biotechne | Cat# AF1157 |
| mouse anti-α-SMA | Biolegend | Cat# 904601 |
| mouse anti-E-cadherin | Cell Signaling | Cat# 14472S |
| mouse anti-CK7 | Abcam | Cat# ab9021 |
| mouse anti-CK18 | Prozen | Cat# 690028S |
| rabbit anti-Mist | Abcam | Cat# ab187978 |
| mouse anti-dsRed | Takara Bio | Cat# 632392 |
| rabbit anti-RFP | Rockland NB46390SS | Cat# RK600-401-379 |
| EPCAM FITC (clone 9C4) | Biolegend | Cat# 324204 |
| NGFR APC (clone ME20.4) | Biolegend | Cat# 345107 |
| CD24 APC (clone ML5) | Biolegend | Cat# 311117 |
| CD49f (ITGA6) APC (clone GoH3) | Biolegend | Cat# 313615 |

